# Genome assembly of *Danaus chrysippus* and comparison with the Monarch *Danaus plexippus*

**DOI:** 10.1101/2021.11.27.470194

**Authors:** Kumar Saurabh Singh, Rishi De-Kayne, Kennedy Saitoti Omufwoko, Dino J. Martins, Chris Bass, Richard ffrench-Constant, Simon H. Martin

**Author notes:** These authors contributed equally (Kumar Saurabh Singh, Rishi De-Kayne).

## Abstract

Milkweed butterflies in the genus *Danaus* are studied in a diverse range of research fields including the neurobiology of migration, biochemistry of plant detoxification, host-parasite interactions, evolution of sex chromosomes, and speciation. We have assembled a nearly chromosomal genome for *Danaus chrysippus* (known as the African Monarch, African Queen, and Plain Tiger) using long read sequencing data. This species is of particular interest for the study of genome structural change and its consequences for evolution. Comparison with the genome of the North American Monarch *Danaus plexippus* reveals generally strong synteny, but highlights three inversion differences. The three chromosomes involved were previously found to carry peaks of intra-specific differentiation in *D. chrysippus* in Africa, suggesting that these inversions may be polymorphic and associated with local adaptation. The *D. chrysippus* genome is over 40% larger than that of *D. plexippus*, and nearly all of the additional ∼100 Megabases of DNA comprises repeats. Future comparative genomic studies within this genus will shed light on the evolution of genome architecture.

## Introduction

The genus *Danaus* is perhaps best known for the iconic Monarch butterfly *Danaus plexippus* and its extraordinary migrations in North America. Genomic studies of the Monarch have shed light on host plant detoxification (Tan *et al*. 2019), sex chromosome evolution (Mongue *et al*. 2017; Gu *et al*. 2019) and the genetic basis of migratory behaviour (Zhan *et al*. 2014). Its relative *Danaus chrysippus* is found throughout Africa, the Mediterranean and south Asia, and is known as the African Monarch, African Queen, and Plain Tiger butterfly in different parts of its range. *D. chrysippus* is emerging as a useful study system in evolutionary genomics. Several subspecies of *D. chrysippus* with distinct warning patterns occupy distinct geographic ranges separated by broad hybrid zones (Smith *et al*. 1997; Lushai *et al*. 2003). Patterns of genetic differentiation suggest a role for chromosomal rearrangements in maintaining these differences (Martin *et al*. 2020). In the east African hybrid zone, a neo-W sex chromosome has emerged in the past few thousand years and is associated with infection by a male-killing endosymbiont *Spiroplasma* (Smith *et al*. 2016; Martin *et al*. 2020). This species therefore has great potential for future research on the evolutionary impacts of genome structural change.

Here we describe the generation of a chromosome-level assembly for *D. chrysippus* based on Pacific Biosciences long read sequencing data. This serves to replace a previous reference genome based on short read sequences and mate-pair libraries, which had low contiguity (N50=0.63Mb, Martin *et al*. 2020). Our new assembly has an N50 of 11.45 Mb. Nineteen of the 30 chromosomes are represented by a single contig, and the remaining eleven by two contigs each. At 354 Mb, this genome is average in size for a butterfly, but about 1.4 times larger than that of *D. plexippus* (∼250 Mb). Comparative analyses indicate that this difference is largely explained by increased repeat content, but *D. chrysippus* also has larger introns, implying that these species have experienced different selection pressures acting on non-essential DNA.

## Materials and Methods

### DNA sequencing

High-molecular-weight DNA was extracted from a single female pupa from a captive butterfly stock using the Qiagen Blood & Cell Culture DNA Mini Kit following the manufacturer’s guidelines. Long read Pacific Biosciences sequencing was performed using seven PacBio Sequel SMRT cells on the Sequel platform, yielding approximately 9.7 gigabases (Gb) per SMRT cell. The 3.8 million PacBio reads totalled 67.6 Gb and had an N50 of 27.3 kb. In addition, we generated Illumina sequencing data for the same individual on the Novaseq 600 platform (118 million paired-end reads of 150 bp with an insert size of 350 bp) totalling 35 Gb.

### Genome assembly

We assembled the long reads using both Canu (Koren *et al*. 2017) and Falcon (Chin *et al*. 2016), and then merged these assemblies to maximize the genome completeness using quickmerge -v 0.3 (Chakraborty *et al*. 2016). Redundant contigs or haplotigs were removed using Purge_haplotigs -v 1.0.4 (Roach *et al*. 2018) with the *-align_cov* (Percent cutoff for identifying a contig as haplotig) value of 65. Before merging, assemblies were polished iteratively using three rounds of Pilon -v 1.22 in diploid mode (Walker *et al*. 2014; using a trimmed version of the short-read data; reads were trimmed using Trim_Galore -v 0.4.0; Krueger 2012), and Racon -v 1.3.1 (Vaser *et al*. 2017; using the long-read data). Illumina and PacBio raw reads are archived under European Nucleotide Archive project accession: PRJEB47812.

### Whole genome alignment and synteny assessment

To assess synteny and putatively assign contigs to chromosomes, we aligned the *D. chrysippus* assembly to two *Danaus plexippus* assemblies: ‘Dplex_v4’, a chromosome-level assembly produced by scaffolding 4115 scaffolds using chromatin conformation (Hi-C) data (GCA_009731565.1, (Gu *et al*. 2019)) and ‘MEX_DaPlex’, a long-read based assembly consisting of 66 scaffolds, of which 38 (97% of total sequence) have been assigned to chromosomes (GCA_018135715.1, (Ranz *et al*. 2021)). Alignments were generated using both MUMmer’s nucmer tool version 3.1 (Marçais *et al*. 2018), with default parameters except the ‘maxGap’ parameter set to 1000, and with minimap2 v2.17 (Li 2018), using the ‘asm20’ parameter preset, designed for whole genome alignment of species with sequence divergence below 20%. Nucmer alignments were explored using the interactive alignment visualisation tool Dot (https://github.com/dnanexus/dot) and final alignment plots based on minimap2 alignments were generated using Asynt (https://github.com/simonhmartin/asynt).

### Correcting putative misassemblies

Visualisation of whole genome alignment to both *Danaus plexippus* assemblies (described above) revealed three putatively misassembled contigs that had portions aligning confidently to two different chromosomes. Although these could theoretically represent real translocation or fusion products, we took the conservative decision to split these contigs, as the rearrangements are not apparent in additional long-read assemblies generated in a related study (unpublished data). Optimal split points were identified by visual inspection of the alignments, as well as additional BLASTn alignments made between the two genomes. The original unsplit assembly, along with details of split points, is available at https://doi.org/10.5281/zenodo.5731560.

Despite having performed automated removal of redundant contigs using Purge_haplotigs, visual exploration of the alignments identified a further four contigs that appeared to be redundant (i.e. representing a part of the genome already represented by a larger contig). These included one of the split products described above. To confirm this, we aligned the Illumina reads for the assembled individual back to the assembly using BWA MEM (Li and Durbin 2010) using default parameters, and computed read depth using Samtools depth (Li *et al*. 2009). Visualisation of median read depth averaged in 50 kb windows confirmed that these four contigs had 50% depth, so they were removed from the assembly. Finally, two contigs included portions that appeared to be redundant in the alignments as well as read depth plots. These were therefore split at the point in the alignment where the redundancy began, and the redundant fragment was removed from the final assembly. The full original assembly, along with details of all splits and portions retained to produce the final Dchry2.2 assembly, is available at https://doi.org/10.5281/zenodo.5731560. To assess the base-level accuracy of our assembly we calculated the consensus quality (QV), comparing the frequency of k-mers present in the raw Illumina reads with those present across the final assembly (all 83 contigs) using Merqury v.1.3 (Rhie *et al*. 2020).

### Repeat Annotation

To assess the repeat content of the assembly, the genome was masked using a custom repeat library. First, a repeat library was produced using the finished genome assembly, using RepeatModeler v2.0.1 (Smit and Hubley 2008), and this library was then combined with a broad Lepidoptera repeat database extracted using RepeatMasker v.4.1.0 (Smit *et al*. 2015). Repeat masking of the genome was then carried out using RepeatMasker (Smit *et al*. 2015). To determine the prevalence of expanding transposable element families within *Danaus chrysippus* the scripts calDivergenceFromAlign.pl and createRepeatLandscape.pl from RepeatModeler (Smit and Hubley 2008) were used to produce a repeat landscape for the assembly. To facilitate a comparison with other *Danaus* genome assemblies, this repeat masking process was repeated using the same custom repeat library for two well-assembled *Danaus plexippus* genome assemblies (NCBI accessions GCF_009731565.1 and GCA_018135715.1 (Ranz *et al*. 2021)). The resulting softmasked assemblies were then used for genome annotation.

### Gene Annotation

Due to a lack of RNAseq data, a preliminary genome annotation was carried out using two protein sets from the close relative to *D. chrysippus, D. plexippus*, the Monarch butterfly. This combined protein set was produced by collating protein information from two different, *D. plexippus* assemblies, the first a proteome downloaded from uniprot under the accession UP000596680 (associated with the Dplex_v4 assembly), and the second taken by extracting amino acid sequences from the annotation of the ‘MEX_DaPlex’ *Danaus plexippus* assembly GCF_009731565.1 (Ranz *et al*. 2021) (both protein sets had high BUSCO scores, indicative of high-quality annotation). This combined protein set was then used as input for the BRAKER2 pipeline (Hoff *et al*. 2018) to annotate each of the three soft masked genome assemblies produced above (specifying --gff3 --softmasking --prot_seq=protein_set.fasta --prg=gth -- gth2traingenes –trainFromGth). GenomeTools (Gremme et al. 2013) was then used to sort and tidy the annotation output (gt gff -sort -tidy -retainids -fixregionboundaries) and calculate summary statistics of the annotation (gt stat -genelengthdistri -genescoredistri -exonlengthdistri -exonnumberdistri -intronlengthdistri -cdslengthdistri). Functional annotation for the resulting *Danaus chrysippus* protein set was carried out using Pannzer2 (Törönen *et al*. 2018). To determine variation in intron and exon length between *D. chrysippus* and *D. plexippus*, introns and exons were extracted from our corresponding annotation file for each of the three assemblies.

### Genome comparison and assembly validation

To assess the quality of annotation, BUSCOs were calculated for both the full *D. chrysippus* assembly and the protein sequences resulting from annotation specifying the insecta_odb10 lineage BUSCO set in BUSCO v.5.0.0 (Simão *et al*. 2015). Additionally, the annotation was compared against those of both published *D. plexippus* annotations. To compare each of the assemblies, and in turn the consistency of genome structure across *Danaus* species we plotted the distribution of intron lengths for annotations from each of the three assemblies. This was carried out by extracting introns and exons from the longest annotated transcript for each gene within each of the annotations (using the BRAKER2 re-annotations for both *D. plexippus* assemblies to ensure lengths were comparable across assemblies).

### Data availability

The assembly and annotation are available at the European Nucleotide Archive project accession: PRJEB47812. Additional data files are provided at https://doi.org/10.5281/zenodo.5731560: purged haplotigs, assembly before manual edits, details of manual edits made to the assembly, and repeat library and functional annotation files. Scripts for genome assembly are available at https://github.com/kumarsaurabh20/DChry2.1 and scripts for the genome annotation and analysis of introns and exons at https://github.com/RishiDeKayne/Danaus_Dchry2.2_annotation.

## Results and Discussion

### Genome assembly

67.6 Gb of long-read data was assembled into 83 contigs. Manual splitting of three putatively misassembled contigs and removal of several remaining redundant fragments (see Materials and Methods) left 83 contigs with an N50 of 11.45 Mb and L50 of thirteen contigs, giving a total genome size of 354 Mb. Alignment with two different *D. plexippus* assemblies allowed us to confidently assign 41 contigs representing 97% of the sequence length to chromosomes (Figure 1). Of the 30 *D. chrysippus* chromosomes, 19 are represented by a single contig and the rest by two contigs each. The contiguity of our assembly is therefore comparable to that of the *D. plexippus* ‘MEX_DaPlex’ assembly (Ranz *et al*. 2021), for which 38 out of 66 scaffolds (97% of the assembly) were assigned to chromosomes, of which 23 are represented by a single scaffold. Among the 42 *D. chrysippus* contigs that were not assigned to a chromosome (3% of the genome) it is possible that some represent fragments of the female-specific W chromosome.

**Figure 1.**
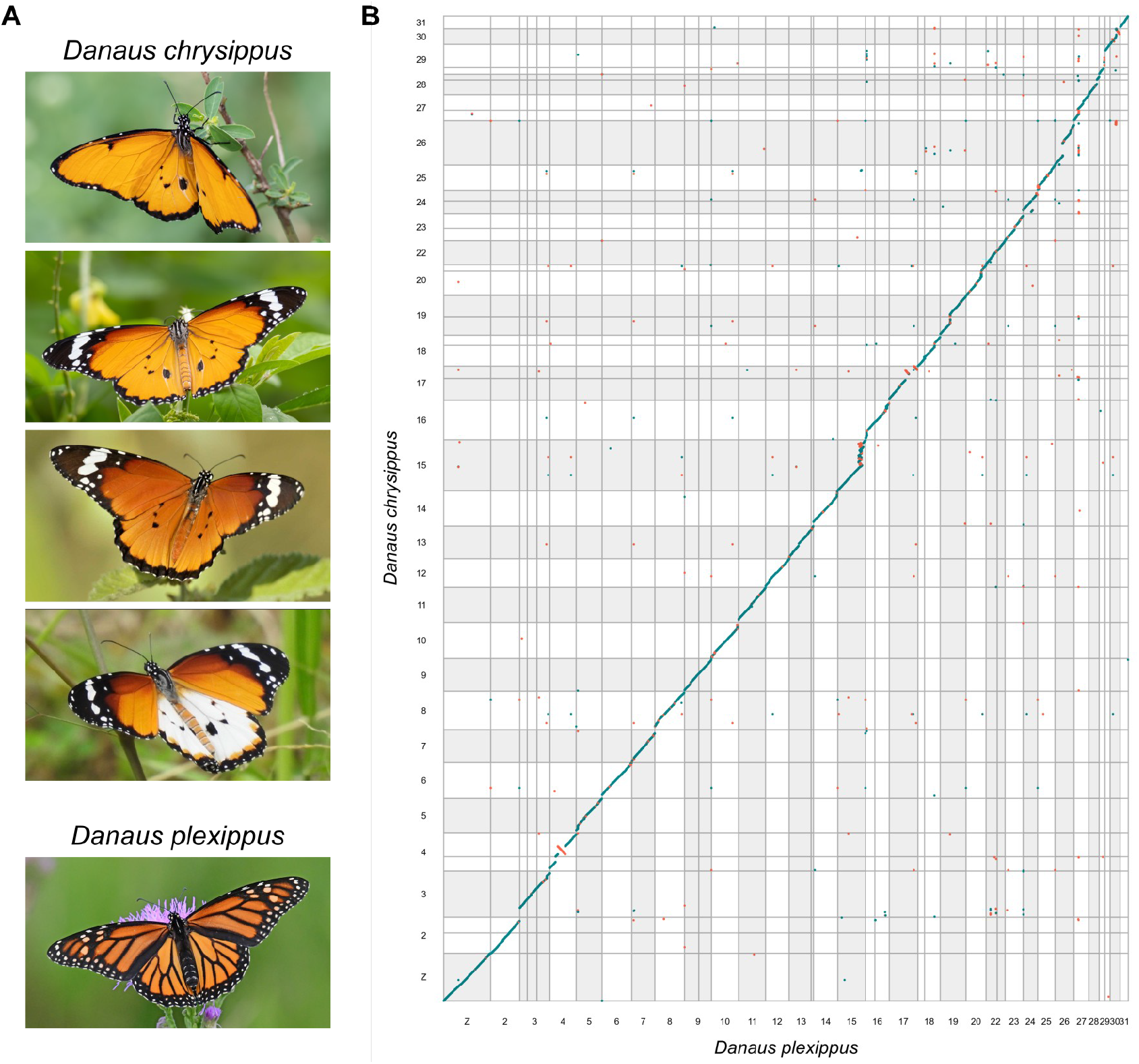
(**A**) Four colour morphs of *D. chrysippus* (above) and *D. plexippus* (below). (**B**) Whole genome alignment between *D. chrysippus* and *D. plexippus* (MEX_DaPlex assembly). Points represent minimap2 alignments greater than 2kb in length. Alignments in the same orientation are shown in turquoise and those in the reverse orientation are shown in red. Only contigs that were confidently assigned to chromosomes (97% of the total in both assemblies) are included. Alternating grey and white bars indicate separate chromosomes. See Figure S1 for the same plot including contig/scaffold labels. Butterfly images from top to bottom by Forest Jarvis (CC-BY-NC), Paul Dickson (CC-BY-NC), Claude Martin, Steven Schulting (CC-BY-NC) and Edward Perry IV (CC-BY-NC).

However, given that butterfly W chromosomes are highly repetitive and have low inter-specific homology (Lewis *et al*. 2021), further work comparing male and female genomes is required to test this hypothesis. The genome-wide consensus quality of the assembly (QV; representing a log-scaled probability of error for each base in our assembly) was 36.2373, suggesting a high level of accuracy (equating to an accuracy between 99.9% and 99.99%).

### Synteny and genome size comparison

The genomes of *D. chrysippus* and *D. plexippus* are largely syntenic (Figure 1). Our assembly supports the earlier finding that the Z sex chromosome of *Danaus* species represents a fusion between the ancestral lepidopteran Z chromosome and autosome 21, which occurred in a recent ancestor of the genus (Mongue *et al*. 2017). We numbered chromosomes according to their homologs in the most recent *D. Plexippus* assembly (Ranz *et al*. 2021), which follows the growing convention of using the chromosome numbering system introduced for *Melitaea cinxia*, the first assembled lepidopteran genome that retains the ancestral karyotype of 31 (Ahola *et al*. 2014; Cicconardi *et al*. 2021; Ranz *et al*. 2021; Lewis *et al*. 2021). As such, the *Danaus* karyotype lacks an autosome 21, as this is now part of the Z sex chromosome.

Several putative inversion differences can be identified between the two *Danaus* species, most notably on chromosomes 4, 17 and 30 (Figure 1). We note that all three of these chromosomes were found to carry sharp peaks of intra-specific differentiation between subspecies of *D. chrysippus* in Africa, against a background of very low genetic differentiation (Martin *et al*. 2020), suggesting that these putative inversions may be polymorphic and associated with local-adaptation in *D. chrysippus*. In addition, chromosomes 15, 26 and 29 all carry large duplicated/repetitive portions relative to *D. plexippus*. One of these, on chromosome 15, was identified previously as the site of a large expansion in gene copy number through multiple duplications and is associated with subspecies differentiation and colour pattern variation in *D. chrysippus* (Martin *et al*. 2020). Further work to dissect this genomic region and compare chromosome structure among *D. chrysippus* subspecies is ongoing.

In total, the 354 Mb *D. chrysippus* genome is 42-44% larger than that of *D. plexippus* (245-248 Mb). This difference is consistent for all autosomes, but most dramatic for the three chromosomes carrying large repetitive/duplicated tracts: namely 15, 26 and 29 (Figure 2). By contrast, the Z sex chromosome is nearly identical in size in the two species. This difference in autosome sizes could be explained either via a systematic size reduction in the lineage leading to *D. plexippus*, or a systematic increase in the lineage leading to *D. chrysippus*. These hypotheses can be distinguished by comparison with assemblies of other members of the genus or outgroups in the future.

**Figure 2.**
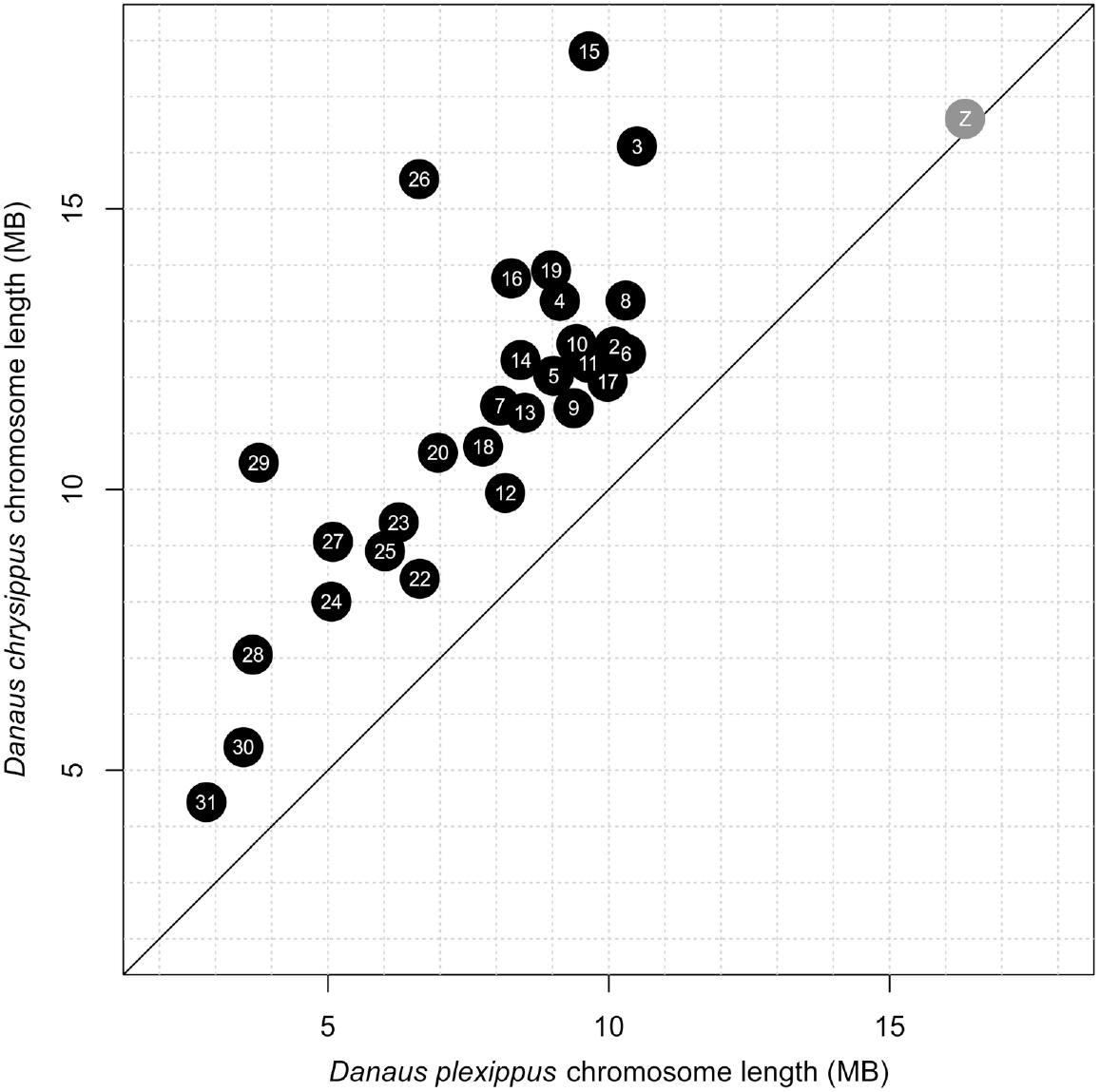
Chromosome length comparison between *D. chrysippus* and *D. plexippus* (MEX_DaPlex assembly). Chromosome lengths represent the sum total of the contigs/scaffolds assigned to each chromosome. Autosomes are shown in black and the Z sex chromosome in grey.

### Transposable element and repeat content

In total the *D. chrysippus* genome comprises 35.5% repeats, with the largest proportion of these being retroelements which make up 11.9% of the genome sequence (Figure 3). Repeat masking of each of the *D. plexippus* assemblies revealed that a substantially lower proportion of the genomes of these close relatives comprise repeats, only between 11.2% and 14.3%. Each of the main classified repeat families are more abundant in *D. chrysippus* compared to *D. plexippus*, with the largest difference between the species observed for the rolling-circle family which represents 1.8% and 2% of the *D. plexippus* genome sequence, compared to 7.6% of the *D. chrysippus* sequence. The repeat landscape of *D. chrysippus* (Figure 4) highlights a number of expanding repeat families, most strikingly the rolling-circle repeats RC/Helitron (pink; Figure 4). The increased prevalence of repetitive elements within the *D. chrysippus* genome (91-98Mb more than *D. plexippus*) largely explains the larger genome size of *D. chrysippus* compared to *D. plexippus* (an increase of 106-109Mb). Although the repeat content of genomes across the Lepidoptera order has been shown to vary substantially (Talla *et al*. 2017) our results suggest that even within a genus a large amount of variation can be present. Although the genome of *D. chrysippus* is rather repetitive, even within Lepidoptera, (With repeats making up 35.5% of the genome), *D. plexippus* tends towards the lower end of repeat content (repeats make up 11.2-14.3% of the genome).

**Figure 3.**
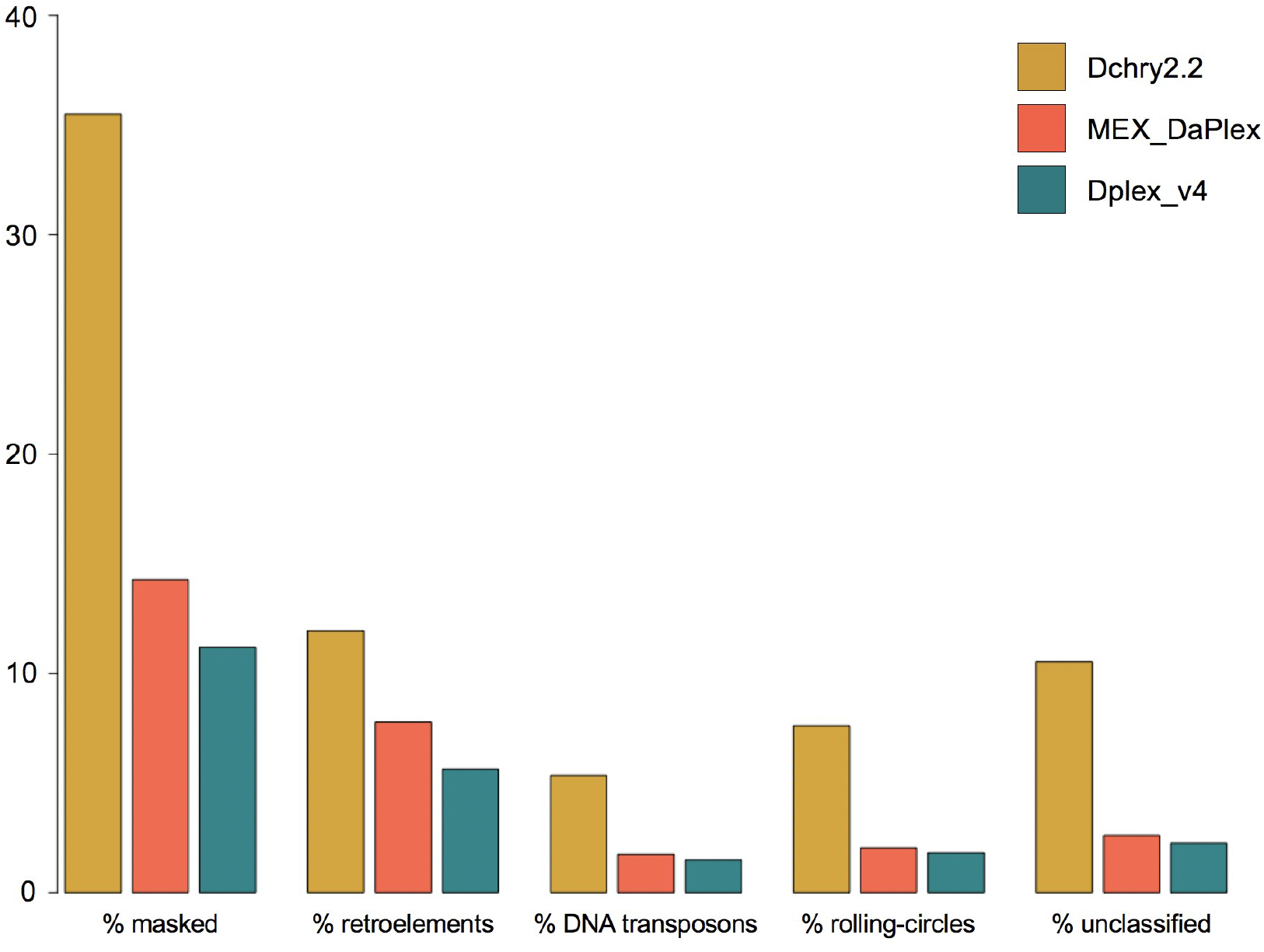
Barplot showing the proportion of the genome of *Danaus chrysippus* (yellow) and *D. plexippus* (represented by both the MEX_DaPlex ,in red, and Dplex_v4, in turquoise, assemblies) made up of repetitive elements (as identified and masked by repeatmodeler and repeatmasker). Additionally, the proportion of each genome made up of specific repetitive element families is shown highlighting the increased proportion of repetitive elements in *Danaus chrysippus*.

**Figure 4.**
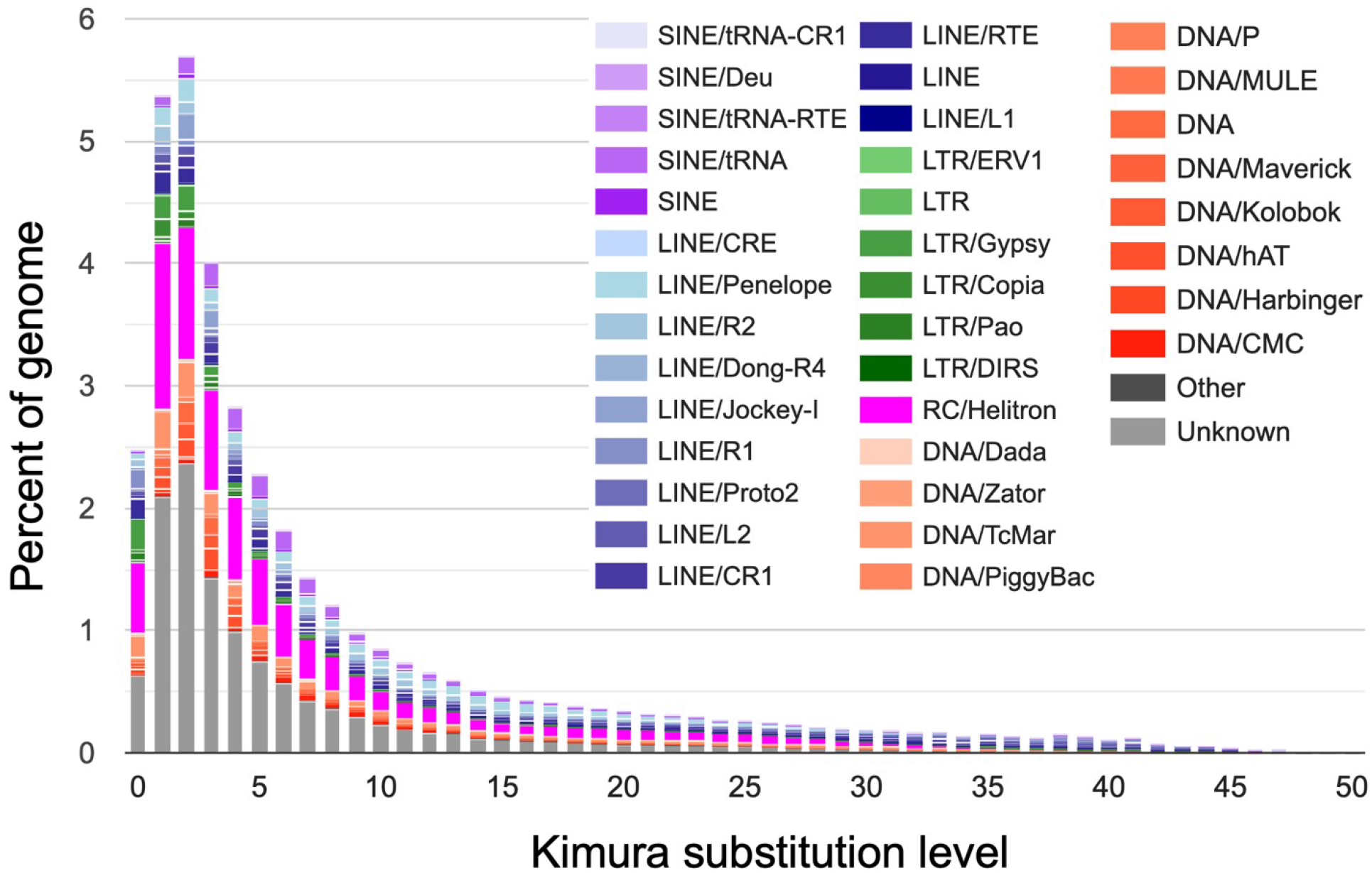
The repeat landscape of the Dchry2.2 assembly. In addition to unclassified repeats, rolling-circle (RC/Helitron), LINE and LTR families all appear to have expanded recently.

**Figure 5.**
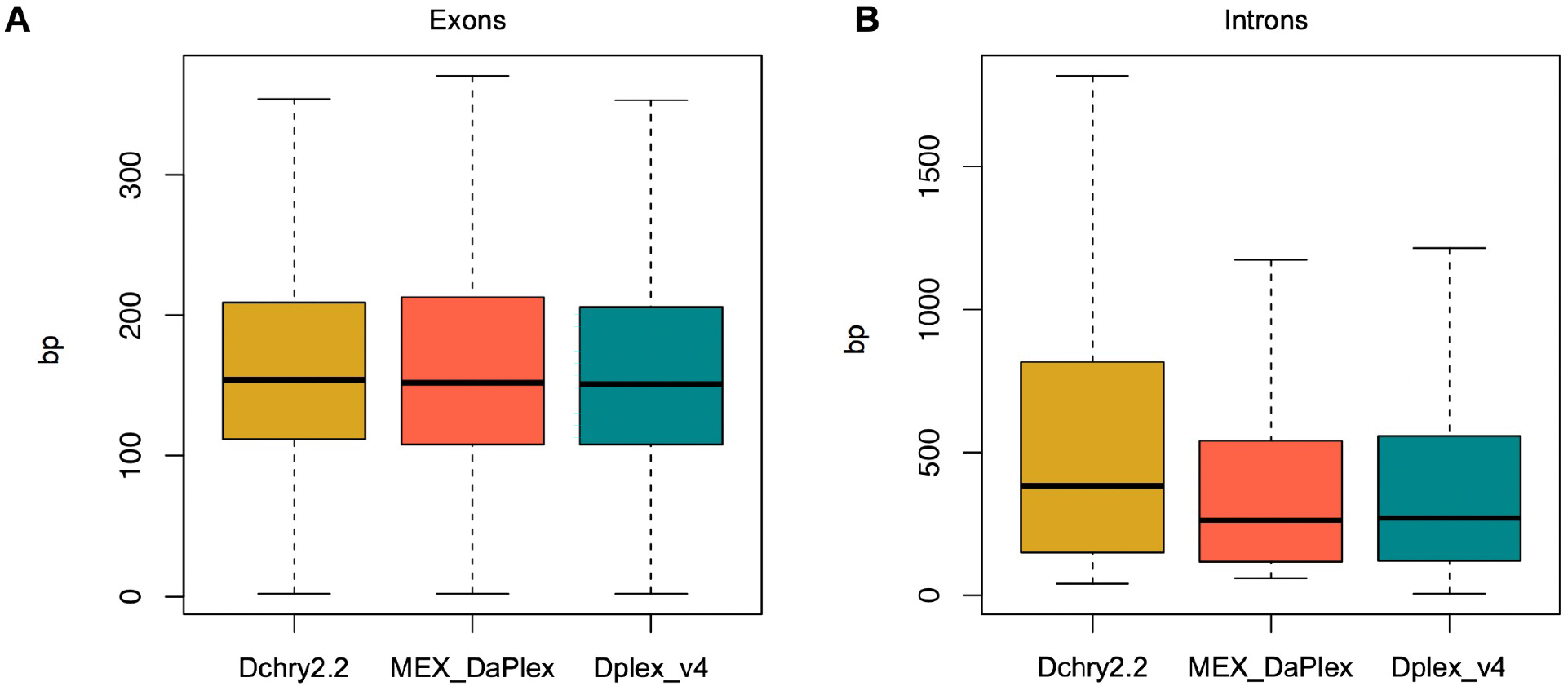
Boxplots showing the distribution of (A) exon lengths and (B) intron lengths taken from the longest transcripts annotated with BRAKER2 for the three assemblies, *D. chrysippus* (Dchry2.2) and each of the re-annotated *D. plexippus* assemblies (MEX_DaPlex and Dplex_v4). Outlier points were omitted for clarity. Mean exon length is 226bp in Dchry2.2, 238bp for the re-annotated MEX_DaPlex assembly, and 217bp for the re-annotated Dplex_v4 assembly. Mean intron length is 975bp in Dchry2.2, 665bp for the re-annotated MEX_DaPlex assembly, and 738bp for the re-annotated Dplex_v4 assembly.

### Gene content

In total 16,260 protein-coding genes were annotated in the assembly by BRAKER2 (with 19,639 protein-coding mRNAs annotated - accounting for multiple transcripts/isoforms of the same gene), which included 136,694 exons and 117,106 introns. This number of genes is similar to that of the published annotations for each of the *D. plexippus* assemblies, which annotated 15,006 (Dplex_v4) and 15,995 (MEX_DaPlex) genes (as well as our re-annotated versions of these assemblies which annotated 18,663 and 21,311 genes; in both cases our annotation involved annotating additional smaller scaffolds not annotated in the original assemblies - 284 vs 30 scaffolds for Dplex_v4 and 64 vs 55 for MEX_DaPlex). An analysis of BUSCOs using the *insecta_odb10* benchmarking set shows that the full genome sequences and annotated protein set for *D. chrysippus* are 98.2% and 96.3% complete for BUSCOs, respectively. This percentage completeness is close to that of both published *D. plexippus* annotations which have 94.6% (Dplex_v4) and 97.1% (MEX_DaPlex) complete BUSCOs (Table 1). Pannzer2 allowed us to add functional annotation to 9,567 of the full 16,260 gene set (functional annotation available at https://doi.org/10.5281/zenodo.5731560).

**Table 1.** BUSCOs for the *D. chrysippus* genome assembly and each *D. plexippus* assembly in addition to the protein sequences resulting from both the original and re-annotation of each of these assemblies (using the insecta_odb10 BUSCO set, n=1367).

|  |  | % complete | % single | % duplicated | % fragmented | % missing |
| --- | --- | --- | --- | --- | --- | --- |
| Genomes | **Dchry2.2** | 98.2 | 97.5 | 0.7 | 0.6 | 1.2 |
|  | Dplex_v4 | 98.7 | 98.2 | 0.5 | 0.7 | 0.6 |
|  | MEX_DaPlex | 98.9 | 98.2 | 0.7 | 0.4 | 0.7 |
| Protein sets | **Dchry2.2** | 96.3 | 79.7 | 16.6 | 1.1 | 2.6 |
|  | Dplex_v4 | 94.6 | 93.7 | 0.9 | 2.5 | 2.9 |
|  | Dplex_v4<br>(re-annotated) | 98.1 | 72.6 | 25.5 | 1.2 | 0.7 |
|  | MEX_DaPlex | 97.1 | 87.1 | 10.0 | 1.3 | 1.6 |
|  | MEX_DaPlex<br>(re-annotated) | 98.8 | 72.6 | 26.2 | 0.7 | 0.5 |

### Intron and exon length

Exon length is relatively consistent across the three genomes (ranging from 217bp - Dplex_v4 re-annotation to 238bp in the MEX_DaPlex re-annotation). However, mean intron length in the *D. chrysippus* is higher than that of the two *D. plexippus* assemblies (ranging from 665bp to 738bp, in the MEX_DaPlex re-annotation and Dplex_v4 re-annotation, respectively), with a mean of 975bp. This substantial increase in intron length within *D. chrysippus* likely explains the remaining variation in genome size between *D. chrysippus* and *D. plexippus*. This difference may represent a neutral increase in introns in *D. chrysippus* or a selection-mediated reduction in intron size in *D. plexippus*. These hypotheses may be resolved by comparison with genomes of other members of the genus in the future.

## Conclusions

We have assembled a nearly chromosome-level genome for *D. chrysippus*, which is highly comparable in its quality to the best available assembly for *D. plexippus*. Although the two species retain strong synteny, the *D. chrysippus* genome is >40% larger, with more repetitive content and larger introns on average. This implies stronger selection to limit non-essential DNA in *D. plexippus*. Future comparative studies involving other members of the genus could shed light on the processes and forces driving the evolution of genome size. The *D. chrysippus* genome will also serve as a reference for population genomic studies to test hypotheses about the evolution of warning colouration, host-parasite interactions and the consequences of chromosomal rearrangements.

## Acknowledgements

We thank Alex Mackintosh for providing advice on genome assembly and annotation.

## Supplementary Figures

**Figure S1.**
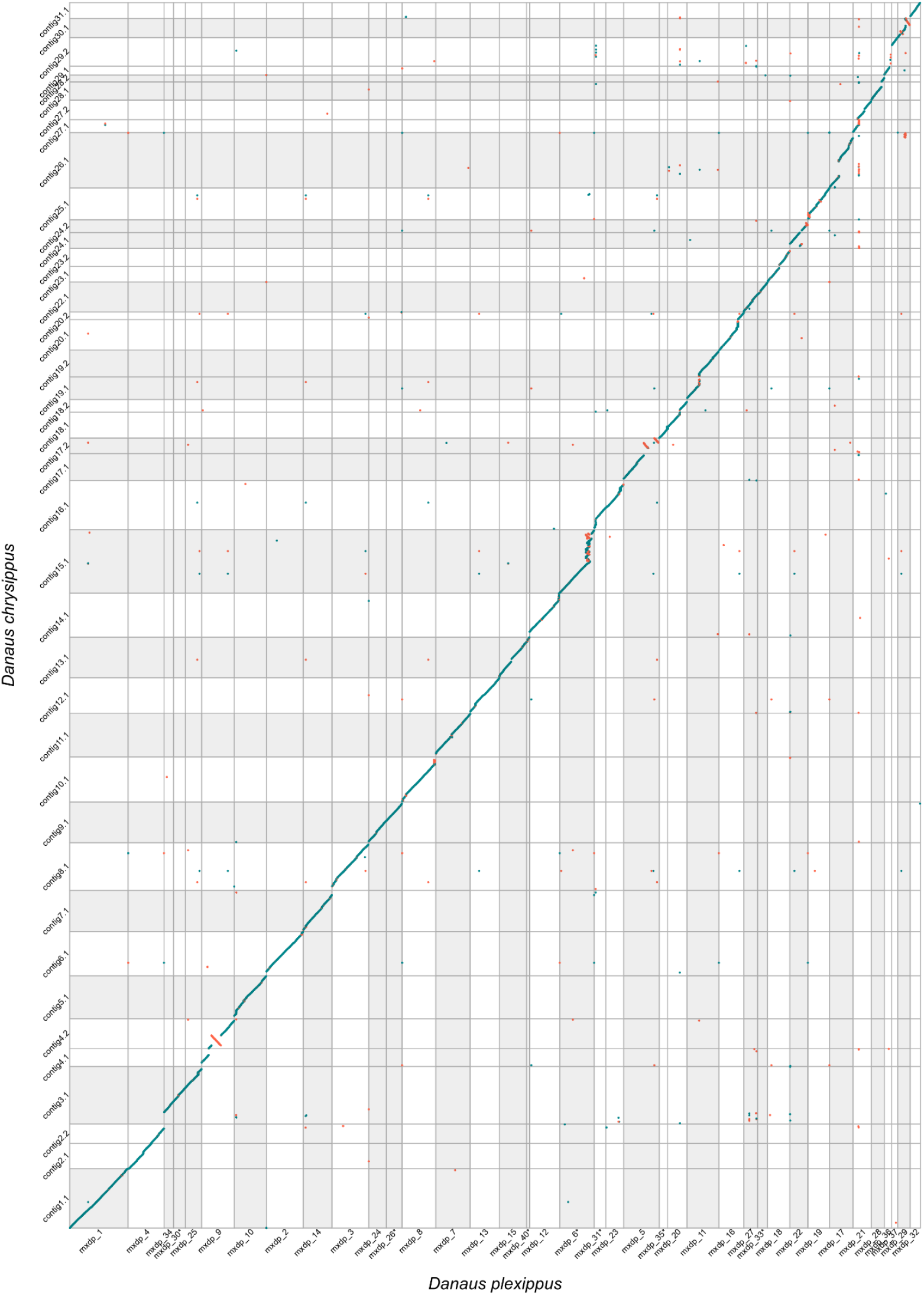
Whole genome alignment between *D. chrysippus* and *D. plexippus* (MEX_DaPlex assembly) showing contig and scaffold labels. Points represent minimap2 alignments greater than 2kb in length. Alignments in the same orientation are shown in turquoise and those in the reverse orientation are shown in red. Only contigs/scaffolds that were confidently assigned to chromosomes (97% of the total in both assemblies) are included. Chromosomes are shaded alternately in grey and white. *D. plexippus* scaffolds that were reversed for ease of visualisation are indicated by an asterisk after the label.

## References

Ahola, V., R. Lehtonen, P. Somervuo, L. Salmela, P. Koskinen et al., 2014 The Glanville fritillary genome retains an ancient karyotype and reveals selective chromosomal fusions in Lepidoptera. Nat. Commun. 5: 4737.

Chakraborty, M., J. G. Baldwin-Brown, A. D. Long, and J. J. Emerson, 2016 Contiguous and accurate de novo assembly of metazoan genomes with modest long read coverage. Nucleic Acids Res. 44: e147.

Chin, C.-S., P. Peluso, F. J. Sedlazeck, M. Nattestad, G. T. Concepcion et al., 2016 Phased diploid genome assembly with single-molecule real-time sequencing. Nat. Methods 13: 1050–1054.

Cicconardi, F., J. J. Lewis, S. H. Martin, R. D. Reed, C. G. Danko et al., 2021 Chromosome fusion affects genetic diversity and evolutionary turnover of functional loci, but consistently depends on chromosome size. Mol. Biol. Evol. 38: 4449–4462

Gremme, G., S. Steinbiss, and S. Kurtz, 2013 GenomeTools: a comprehensive software library for efficient processing of structured genome annotations. IEEE/ACM Trans. Comput. Biol. Bioinform. 10: 645–656.

Gu, L., P. F. Reilly, J. J. Lewis, R. D. Reed, P. Andolfatto et al., 2019 Dichotomy of Dosage Compensation along the Neo Z Chromosome of the Monarch Butterfly. Curr. Biol. 29: 4071–4077.

Hoff, K. J., A. Lomsadze, M. Stanke, and M. Borodovsky, 2018 BRAKER2: incorporating protein homology information into gene prediction with GeneMark-EP and AUGUSTUS. Plant and Animal Genomes XXVI.

Kolmogorov, M., J. Yuan, Y. Lin, and P. A. Pevzner, 2019 Assembly of long, error-prone reads using repeat graphs. Nat. Biotechnol. 37: 540–546.

Koren, S., B. P. Walenz, K. Berlin, J. R. Miller, N. H. Bergman et al., 2017 Canu: scalable and accurate long-read assembly via adaptive k-mer weighting and repeat separation. Genome Res. 27: 722–736.

Krueger, F., 2012 Trim Galore: a wrapper tool around Cutadapt and FastQC to consistently apply quality and adapter trimming to FastQ files, with some extra functionality for MspI-digested RRBS-type (Reduced Representation Bisufite-Seq) libraries. URL http://www.bioinformatics.babraham.ac.uk/projects/trim_galore/. (Date of access: 28/04/2016).

Lewis, J. J., F. Cicconardi, S. H. Martin, R. D. Reed, C. G. Danko et al., 2021 The Dryas iulia Genome Supports Multiple Gains of a W Chromosome from a B Chromosome in Butterflies. Genome Biol. Evol. 13: evab128

Li, H., 2018 Minimap2: pairwise alignment for nucleotide sequences. Bioinformatics 34: 3094–3100.

Li, H., and R. Durbin, 2010 Fast and accurate long-read alignment with Burrows-Wheeler transform. Bioinformatics 26: 589–595.

Li, H., B. Handsaker, A. Wysoker, T. Fennell, J. Ruan et al., 2009 The Sequence Alignment/Map format and SAMtools. Bioinformatics 25: 2078–2079.

Lushai, G., D. A. S. Smith, I. J. Gordon, D. Goulson, J. A. Allen et al., 2003 Incomplete sexual isolation in sympatry between subspecies of the butterfly Danaus chrysippus (L.) and the creation of a hybrid zone. Heredity 90: 236–246.

Marçais, G., A. L. Delcher, A. M. Phillippy, R. Coston, S. L. Salzberg et al., 2018 MUMmer4: A fast and versatile genome alignment system. PLoS Comput. Biol. 14: e1005944.

Martin, S. H., K. S. Singh, I. J. Gordon, K. S. Omufwoko, S. Collins et al., 2020 Whole-chromosome hitchhiking driven by a male-killing endosymbiont. PLoS Biol. 18: e3000610.

Mongue, A. J., P. Nguyen, A. Voleníková, and J. R. Walters, 2017 Neo-sex Chromosomes in the Monarch Butterfly, Danaus plexippus. G3 7: 3281–3294.

Ranz, J. M., P. M. González, B. D. Clifton, N. O. Nazario-Yepiz, P.L. Hernández-Cervantes et al., 2021 A de novo transcriptional atlas in Danaus plexippus reveals variability in dosage compensation across tissues. Commun Biol 4: 791.

Rhie, A., B. P. Walenz, S. Koren, & A. M. Phillippy, 2020 Merqury: reference-free quality, completeness, and phasing assessment for genome assemblies. Genome biology, 21: 1–27.

Roach, M. J., S. A. Schmidt, and A. R. Borneman, 2018 Purge Haplotigs: allelic contig reassignment for third-gen diploid genome assemblies. BMC Bioinformatics 19: 460.

Ruan, J., and H. Li, 2019 Fast and accurate long-read assembly with wtdbg2. bioRxiv 530972.

Simão, F.A., R.M. Waterhouse, P. Ioannidis, E.V. Kriventseva, and E.M. Zdobnov, 2015 BUSCO: assessing genome assembly and annotation completeness with single-copy orthologs. Bioinformatics 31: 3210–3212.

Smith, D. A. S., I. J. Gordon, W. Traut, J. Herren, S. Collins et al., 2016 A neo-W chromosome in a tropical butterfly links colour pattern, male-killing, and speciation. Proc. Biol. Sci. 283: 20160821

Smith, D. A. S., D. F. Owen, I. J. Gordon, and N. K. Lowis, 1997 The butterfly Danaus chrysippus (L.) in East Africa: polymorphism and morph-ratio clines within a complex, extensive and dynamic hybrid zone. Zool. J. Linn. Soc. 120: 51–78.

Smit, A. F. A., and R. Hubley, 2008 RepeatModeler Open-1.0. Available fom http://www.repeatmasker.org.

Smit, A. F. A., R. Hubley, and P. Green, 2015 RepeatMasker Open-4.0. 2013--2015.

Talla, V., A. Suh, F. Kalsoom, V. Dinca, R. Vila et al., 2017 Rapid Increase in Genome Size as a Consequence of Transposable Element Hyperactivity in Wood-White (Leptidea) Butterflies. Genome Biol. Evol. 9: 2491–2505.

Tan, W.-H., T. Acevedo, E. V. Harris, T. Y. Alcaide, J. R. Walters et al., 2019 Transcriptomics of monarch butterflies (Danaus plexippus) reveals that toxic host plants alter expression of detoxification genes and down-regulate a small number of immune genes. Mol. Ecol. 28: 4845–4863.

Törönen, P., A. Medlar, and L. Holm, 2018 PANNZER2: a rapid functional annotation web server. Nucleic Acids Res. 46: W84–W88.

Vaser, R., I. Sović, N. Nagarajan, and M. Šikić, 2017 Fast and accurate de novo genome assembly from long uncorrected reads. Genome Res. 27: 737–746.

Walker, B. J., T. Abeel, T. Shea, M. Priest, A. Abouelliel et al., 2014 Pilon: an integrated tool for comprehensive microbial variant detection and genome assembly improvement. PLoS One 9: e112963.

Zhan, S., W. Zhang, K. Niitepõld, J. Hsu, J. F. Haeger et al., 2014 The genetics of monarch butterfly migration and warning colouration. Nature 514: 317–321.

